# Convergent and divergent patterns of holobiont evolution in co-distributed chemosynthetic symbioses

**DOI:** 10.64898/2026.10.01.753749

**Authors:** Corinna Breusing, Caitlin Q. Plowman, Shawn M. Arellano, Andrew J. Davies, Brennan T. Phillips, Craig M. Young, Roxanne A. Beinart

**Affiliations:** Graduate School of Oceanography, University of Rhode Island, Narragansett, RI, USA; Oregon Institute of Marine Biology, University of Oregon, Coos Bay, OR, USA; Department of Biology, University of Oregon, Eugene, OR, USA; Shannon Point Marine Center, Western Washington University, Anacortes, WA, USA; Department of Biology, Western Washington University, Bellingham, WA, USA; Marine and Coastal Science Program, Western Washington University, Bellingham, WA, USA; Department of Biological Sciences, University of Rhode Island, Kingston, RI, USA; Department of Ocean Engineering, University of Rhode Island, Narragansett, RI, USA

**Keywords:** Chemosynthetic symbiosis, cold seeps, host-symbiont population genetics, comparative genomics, convergent evolution, divergent evolution

## Abstract

Intraspecific variation in microbial symbioses is shaped by a variety of ecological and evolutionary processes. In geographically separated populations, divergence in genomic traits is expected due to dispersal barriers and habitat differentiation, whereas sympatric populations might exhibit functional convergence to the characteristics of a shared environment. While these two processes have been well documented in many animal and plant species, they remain understudied in symbiotic systems. Here, we examined these processes in Atlantic cold seep mussel species that associate with environmentally acquired methane and/or sulfur-oxidizing bacteria. Using genomic methods, we reveal an eco-geographical barrier between the Western Atlantic and Gulf of Mexico that influences the distribution of symbiont species and the level of divergence between holobiont populations. While host populations were only slightly differentiated across this divide, symbiont populations were structured both between and within regions. The sulfur-oxidizing symbionts contained different regional types of dissimilatory nitrate reductase, possibly due to variation in oxygen availability. The methane-oxidizing symbionts showed evidence for convergent evolution, such as retention of a secondary, iron-dependent methane monooxygenase in Western Atlantic populations, driven by local copper limitation. These findings advance our understanding of divergent and convergent adaptive mechanisms underlying intraspecific host-symbiont variation in horizontally transmitted symbioses.

**Lay Summary:** Mussels of the subfamily Bathymodiolinae are foundation fauna at deep-sea cold seeps, where they rely on methane- and sulfur-utilizing bacteria for nutrition. Little is known about how these symbioses have evolved across large geographic ranges and varying environmental conditions, although it is thought that mussels typically partner with symbionts whose traits are suited for the specific characteristics at their local habitat. In this study, we collected samples of three co-occurring cold-seep mussel species across thousands of kilometers in the Western Atlantic and Gulf of Mexico and used genomic methods to compare both the mussel hosts and their symbiotic bacteria. Within each species, we found that, while mussel populations did not vary much genetically across the two regions, the genomic traits of the symbiotic bacteria differed between locations. For example, different symbiont species in the Western Atlantic independently shared a comparatively rare enzyme for methane utilization, likely because of the specific environmental conditions at seeps in this region. This observation suggests the occurrence of parallel adaptations across symbiont species, underscoring the importance of environmental pressures in driving the evolution of microbial symbioses.

## Introduction

It is now widely recognized that most animal species live in symbiosis with beneficial microbes that fundamentally impact host evolutionary ecology^1–5^. The function, evolution and persistence of these holobionts is profoundly shaped by symbiont transmission mode. For marine symbiotic animals, horizontal transmission of microbial symbionts is common, even when host animals obligately depend on their symbiont for survival^6,7^. Horizontal transmission carries risks: offspring may fail to obtain their essential partners, or they may be invaded by cheaters or pathogens during the acquisition process^8^. However, these risks may be offset by the fitness advantage of allowing offspring to establish partnerships with locally adapted symbiont strains rather than inheriting potentially maladapted symbionts from their parents. This plasticity becomes especially valuable for hosts whose offspring disperse into new environments that differ substantially from their natal habitat. Indeed, local adaptation via divergent selection in symbiont populations across a host species’ geographic range has been reported in several systems^9,10^.

Even stronger evidence for local adaptation would come from observations of parallel evolution across independent species that occupy similar habitats^11^. Convergent local adaptation has been documented in several taxa, including sympatric but distantly related tree^12^ and woodpecker species^13^, and closely related bee species^14^. Thus, for sympatric horizontally transmitted symbioses, we would predict functional similarities in the symbionts of different host species if local adaptation drives symbiont population structure. Conversely, when organisms share habitats, competition can drive divergence through niche partitioning. We, therefore, might also expect symbionts associated with co-occurring host species to diverge functionally, ultimately enabling species co-existence by driving the holobionts to occupy distinct niches^15–17^. Consequently, intraspecific variation in the symbionts of co-occurring hosts likely results from both evolutionary convergence and divergence, although these processes remain poorly investigated in environmentally acquired symbioses.

Deep-sea mussels of the genera *Bathymodiolus* and *Gigantidas* offer a comparative framework to study divergent and convergent evolution in horizontally transmitted symbioses. In the Atlantic Ocean, three bathymodiolin mussel species (*B. brooksi*, *B. heckerae, G. childressi*) are regionally and sometimes locally sympatric at cold seeps along the Western Atlantic margin (WAM) and Gulf of Mexico (GoM)^18^. These mussels form mutualistic symbioses with environmentally acquired, chemosynthetic bacteria that provide the primary nutrition for their hosts via the oxidation of methane or sulfide emitted in seeping fluid^19–21^. While *G. childressi* associates with a single species of methanotrophic endosymbionts, *B. brooksi* and *B. heckerae* contain both methane-oxidizing (MOX) and sulfur-oxidizing (SOX) bacteria, with *B. heckerae* often harboring multiple SOX species and additional methylotrophic and hydrocarbon-degrading symbiont phylotypes^19,22–25^.

Here we applied genomic methods to determine divergent and convergent processes shaping population structure in *G. childressi*, *B. brooksi* and *B. heckerae* holobionts from the WAM and GoM (Fig. 1A; Table S1). Though some of these species host additional symbionts with potential supplemental functions^19,22–24^, we only focused on the primary SOX and MOX bacteria to enable more direct comparisons among holobionts characterized by single (*G. childressi*), dual (*B. brooksi*) and multiple (*B. heckerae*) symbioses. To evaluate the potential impact of host specificity on symbiont genetic partitioning we also investigated host population differentiation for comparison.

**Fig. 1.**
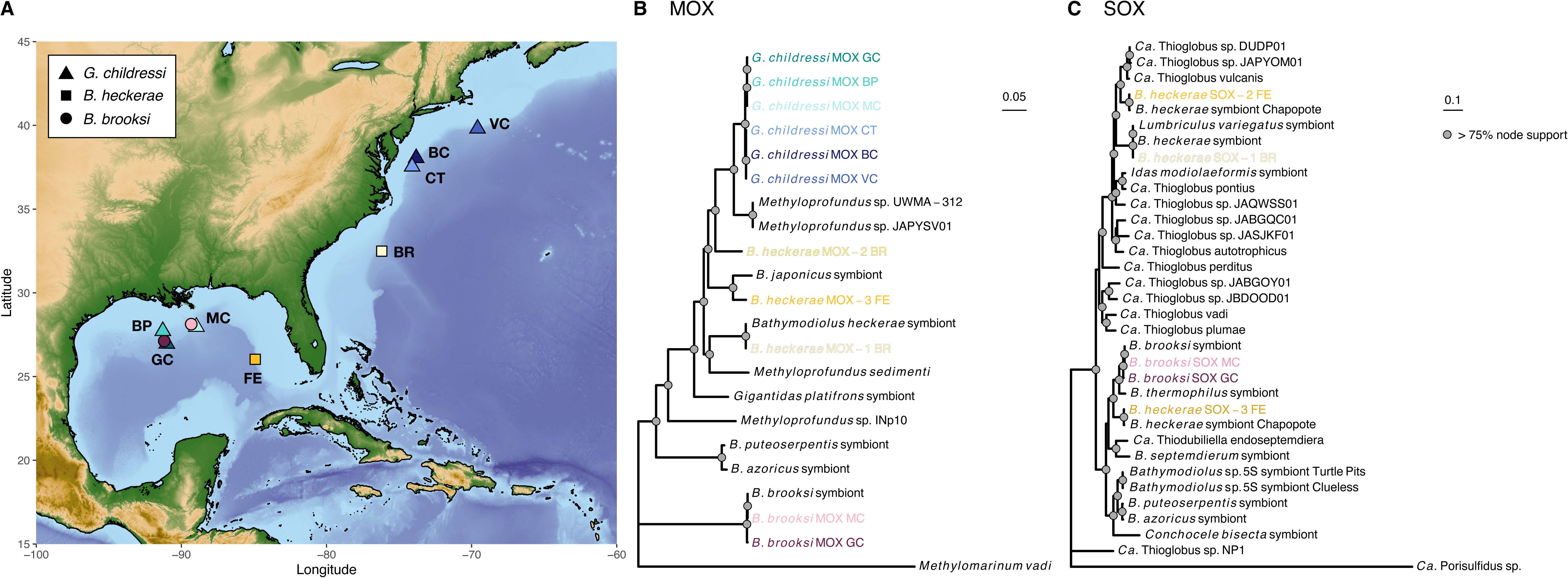
Map of cold seep sampling locations for *G. childressi*, *B. heckerae* and *B. brooksi* in the Western Atlantic and Gulf of Mexico (A) and phylogenetic trees of methane- and sulfur-oxidizing bacteria (B, C). Trees are IQ-Tree consensus phylogenies based on 297 and 33 single-copy core genes, respectively. Grey dots indicate > 75% node support, while scale bars show the expected number of amino acid substitutions per site. *Ca.* Porisulfidus and *Methylomarinum vadi* were used as outgroups to root the Thioglobaceae and Methylomonadaceae specific trees, respectively. VC = Veatch Canyon, BC = Baltimore Canyon, CT = Chincoteague, BR = Blake Ridge, FE = Florida Escarpment, MC = Mississippi Canyon, BP = Brine Pool, GC = GC852.

## Results

### Seep mussels host species-specific, phylogenetically divergent symbionts

Metagenomic binning recovered 223 high-quality symbiont MAGs across mussel samples (MOX = 183; SOX = 40) (Table S2), revealing the presence of five host-specific MOX species (genus *Methyloprofundus*) and four host-specific SOX species (genus *Ca*. Thiodubiliella or *Ca*. Thioglobus). *G. childressi* and *B. brooksi* each hosted a distinct species-specific MOX species across their sampling ranges (Table S2). Three other MOX species occurred solely in *B. heckerae* in a location-specific pattern, with one majority (MOX-1) and one minority (MOX-2) species being observed at Blake Ridge (WAM) and a third species (MOX-3) being observed at Florida Escarpment (GoM) (Table S2). The *B. brooksi* MOX symbiont represents a distantly related *Methyloprofundus* species (Fig. 1B), while the *G. childressi* and *B. heckerae* MOX symbionts belong to a recently diverged clade containing other seep-associated *Methyloprofundus* symbionts and free-living relatives from hydrothermal vents and marine sediments (Fig. 1B). The *G. childressi* MOX symbiont constitutes a well-supported clade with *Methyloprofundus* spp. from the Guaymas Basin to which the *B. heckerae* MOX-2 symbiont is an immediate outgroup. The *B. heckerae* MOX-3 and MOX-1 symbionts form successive sister clades and are placed in proximity to the *B. japonicus* symbiont and *Methyloprofundus sedimenti*, respectively (Fig. 1B).

Sulfur-oxidizing symbionts were only consistently detected in *B. brooksi* and *B. heckerae* (Table S2). We obtained two high-quality MAGs of Arcobacteraceae for *G. childressi* from Veatch Canyon that might be sulfur-oxidizing epibionts^22^. However, due to the rarity of these MAGs we did not further consider them in the analysis. *Bathymodiolus brooksi* contained a single species of *Ca.* Thiodubiliella sp900128405 across seep sites, while *B. heckerae* contained three different, location-specific SOX species (Table S2). One of the *B. heckerae* SOX symbionts, *Ca.* Thioglobus sp. (SOX-1), was observed at Blake Ridge, while the two others, *Ca.* Thioglobus sp900128515 (SOX-2) and *Ca.* Thiodubiliella sp900180345 (SOX-3), which are known from *B. heckerae* at the Chapopote seep in the GoM, were present at Florida Escarpment (Table S2). The *B. brooksi* and *B. heckerae Ca*. Thiodubiliella species are most closely related to the *B. thermophilus* symbiont from Eastern Pacific hydrothermal vents (Fig. 1C), while the two *B. heckerae Ca.* Thioglobus symbionts belong to a broader clade including *Ca.* Thioglobus vulcanis, *Ca.* Thioglobus sp. DUDP01, *Ca.* Thioglobus sp. JAPYOM01 and the symbiont of the annelid *Lumbriculus variegatus* (Fig. 1C).

### Host populations show varying degrees of population structure

The three mussel species differed in population genetic variation based on 36,975–175,062 genomic SNPs (Fig. 2). *Bathymodiolus brooksi* was only collected within the GoM and was virtually undifferentiated between Mississippi Canyon and GC852 (Fig. 2; Table S3; F_ST_ = 0.0032). By contrast, *G. childressi* and *B. heckerae* were obtained from both the WAM and GoM and exhibited population structure between these regions (Fig. 2). *Gigantidas childressi* was slightly differentiated between WAM and GoM localities, with F_ST_s among these regions ranging from 0.0243–0.0290 based on linked SNPs (Table S3; F_ST_ = 0.0181 for pooled data among regions). Results based on unlinked SNPs were almost identical, with pairwise F_ST_s of 0.0252–0.0289 and PCA plots showing comparable levels of separation along axes (Fig. S1; Table S3; F_ST_ = 0.0173 for pooled data among regions). Population differentiation within regions was negligible (F_ST_ < 0.0045; Table S3). Population structure was most pronounced in *B. heckerae*, with a nuclear F_ST_ of 0.0650 (Table S3) and two distinct mitochondrial haplogroups differentiating the Blake Ridge and Florida Escarpment sites (Fig. 2). By contrast, mitochondrial haplotypes in *B. brooksi* and *G. childressi* were not partitioned by geography (Fig. S2, S3).

**Fig. 2.**
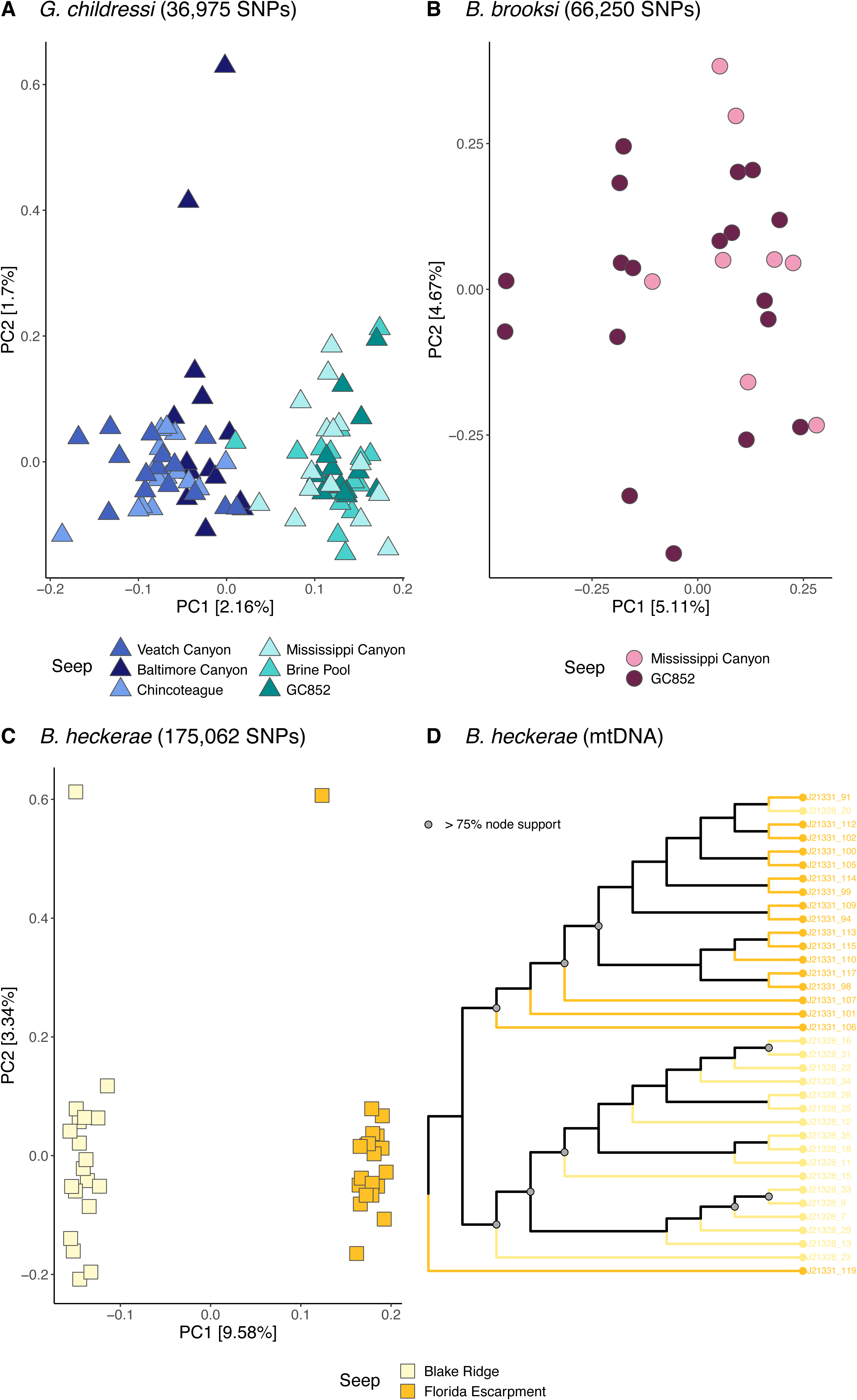
Principal component plots for (A) *G. childressi*, (B) *B. brooksi* and (C) *B. heckerae* host populations based on genomic SNPs. *G. childressi* and *B. heckerae* showed observable population structure between oceanic regions (Western Atlantic versus Gulf of Mexico), while within-region genetic partitioning was limited for all species. In *B. heckerae* notable regional population subdivision was also evident based on mitochondrial haplotypes (D). The cladogram in panel (D) was constructed using complete mitochondrial genomes. See Figs. S2 and S3 for comparison to *G. childressi* and *B. brooksi*.

### Outlier SNPs in host genes are associated with diverse physiological processes

F_ST_ outliers were analyzed in *B. heckerae* and *G. childressi* given observable population structure. In *G. childressi*, 148 outlier SNPs were identified between the WAM and GoM populations, with locus-specific F_ST_s ranging from 0.0877–0.1991 (Table S4). In *B. heckerae*, 734 outlier SNPs were detected, with locus-specific F_ST_s of 0.3897–0.7770 (Table S4).

38 and 221 of the outlier SNPs lay within annotated genes of the *G. childressi* and *B. heckerae* reference genomes, respectively (Table S4, S5). The corresponding genes spanned various physiological functions, including cellular regulation and signaling, RNA and DNA metabolism, membrane transport, cell cycle, immunity and development (Table S5). In both host species some genes were linked to immune system functions and reproductive processes. In *B. heckerae* these included genes for pattern recognition (CD36, fucose-binding protein, ficolin-2-like), inflammatory response modulation (CD200 receptor 1 like, CD109 antigen-like, CD63 antigen-like, myeloid-specific peroxidase), germ cell development (deleted in azoospermia like, septin12) and embryogenesis (distal-less homeobox 6, protein mab-21-like 2) (Table S5). In *G. childressi*, one pattern recognition receptor (MHC class II-related) and two embryogenesis-related genes (NK2 homeobox 4a, cyclin J) were observed (Table S5).

### Symbiont strains are partitioned across geographic regions

Symbiont species were comprised of 2–7 MOX strains and 3–13 SOX strains that were geographically partitioned (Fig. 3). In *G. childressi*, seven MOX strains were strongly differentiated between the WAM and GoM, though some population subdivision was also observed within these regions (Fig. 3A, Fig. S4). A single strain dominated across all WAM seeps, with a second strain being primarily found at Veatch Canyon and Chincoteague, and two other strains being observed at Baltimore Canyon. Three strains were abundant at the GoM seeps, with one strain each being dominant at GC852, Brine Pool and Mississippi Canyon. Pairwise F_ST_ values across regions ranged from 0.8616–0.9134 (0.8442 for data pooled by region), while F_ST_ values within regions ranged from 0.1056–0.3442 without a strict pattern of isolation-by-distance (Table S3). 177 variants represented outlier loci that showed fixed differences among the WAM and GoM populations (Table S6). Most genes containing these variants were poorly characterized based on COG category assignments, followed by functions related to DNA, cell envelope and amino acid metabolism, energy production and inorganic ion transport (Table S6).

**Fig. 3.**
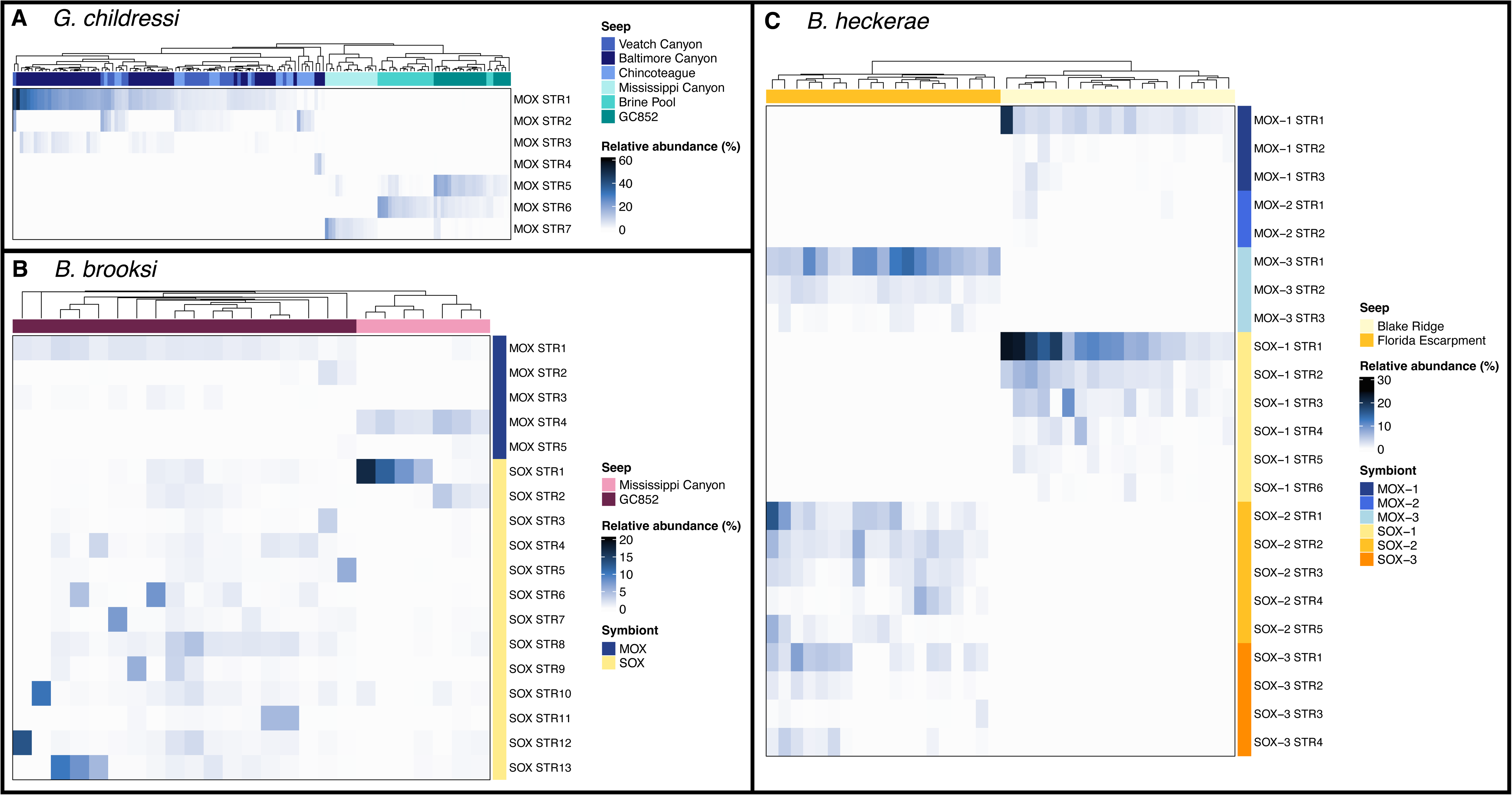
Heatmap of strain relative abundances inferred through StrainFinder for chemosynthetic symbionts of (A) *G. childressi*, (B) *B. brooksi* and (C) *B. heckerae*. In accordance with assessments of symbiont population structure (Fig. S4) and symbiont species distributions, symbiont strains showed both regional and local partitioning between seep sites. Across symbiont species, strain diversity was overall higher in the SOX symbionts than in the MOX symbionts.

Within the *B. brooksi* MOX species, five strains were detected (Fig. 3B). Three strains were primarily observed at GC852, whereas two other strains were mainly present at Mississippi Canyon, with each site harboring one dominant strain. The *B. brooksi* SOX symbiont was comprised of 13 strains, two of which were most prevalent at Mississippi Canyon, while the other eleven SOX strains were primarily observed at GC852 though no single strain appeared to dominate across all host individuals (Fig. 3B). Compared to the MOX strains, the SOX strains showed more overlap between seeps sites, supporting the lower level of population structure observed for this symbiont species (SOX: F_ST_ = 0.2390; MOX: F_ST_ = 0.8974; Fig. S4; Table S3). 1,143 outlier loci exhibited complete differentiation among *B. brooksi* MOX populations, with corresponding gene functions frequently being linked to DNA and energy metabolism, signal transduction, translation and cell envelope metabolism (Table S6). In the *B. brooksi* SOX symbiont only 89 loci were flagged as F_ST_ outliers (F_ST_ = 0.88–1.00). As in the MOX symbiont species, however, these variants were often associated with genes related to replication, recombination and repair as well as cell wall/membrane/envelope biogenesis (Table S6).

In *B. heckerae*, 2–3 strains each were detected for its three MOX species (MOX-1: 3, MOX-2: 2, MOX-3: 3) and 4–6 strains each were detected for its three SOX species (SOX-1: 6, SOX-2: 5, SOX-3: 4) (Fig. 3C). Strains of MOX-1, MOX-2 and SOX-1 were only observed at Blake Ridge, whereas strains of MOX-3, SOX-2 and SOX-3 were only present at Florida Escarpment, with 1–2 strains of each species typically being dominant at the respective sites. Since symbiont species changed between locations, no assessments of symbiont population-level differentiation could be performed for this host species.

### Symbiont species show functional convergence and divergence across geographic regions

The analysis of KOfam presence/absence and enrichment revealed several metabolic distinctions and similarities among the MOX and SOX symbiont species (see Supplementary Results for details). The SOX symbionts comprised three main clusters, consisting of (i) *B. brooksi*’s *Ca.* Thiodubiliella symbiont and *B. heckerae*’s *Ca.* Thiodubiliella SOX-3 (GoM), (ii) *B. heckerae*’s *Ca*. Thioglobus SOX-2 (GoM) and (iii) *B. heckerae*’s *Ca*. Thioglobus SOX-1 (WAM) (Fig. 4A). The cluster of *Ca.* Thioglobus SOX-1 was functionally most divergent from all other, GoM-specific symbiont clusters, regardless of whether the symbionts belonged to the same or different genus (Fig. 4A). The *Ca.* Thiodubiliella symbionts were enriched in multiple proteins for copper detoxification and represented the only taxa with the potential to perform complete denitrification using a membrane-bound *Nar*-type nitrate reductase (Table S7, S8). By contrast, the *Ca.* Thioglobus symbionts appeared to be incomplete denitrifiers, with *B. heckerae*’s *Ca.* Thioglobus SOX-1 symbiont containing a distinct periplasmic *Nap*-type nitrate reductase that was not present in SOX symbionts from the GoM (Table S7, S8).

**Fig. 4.**
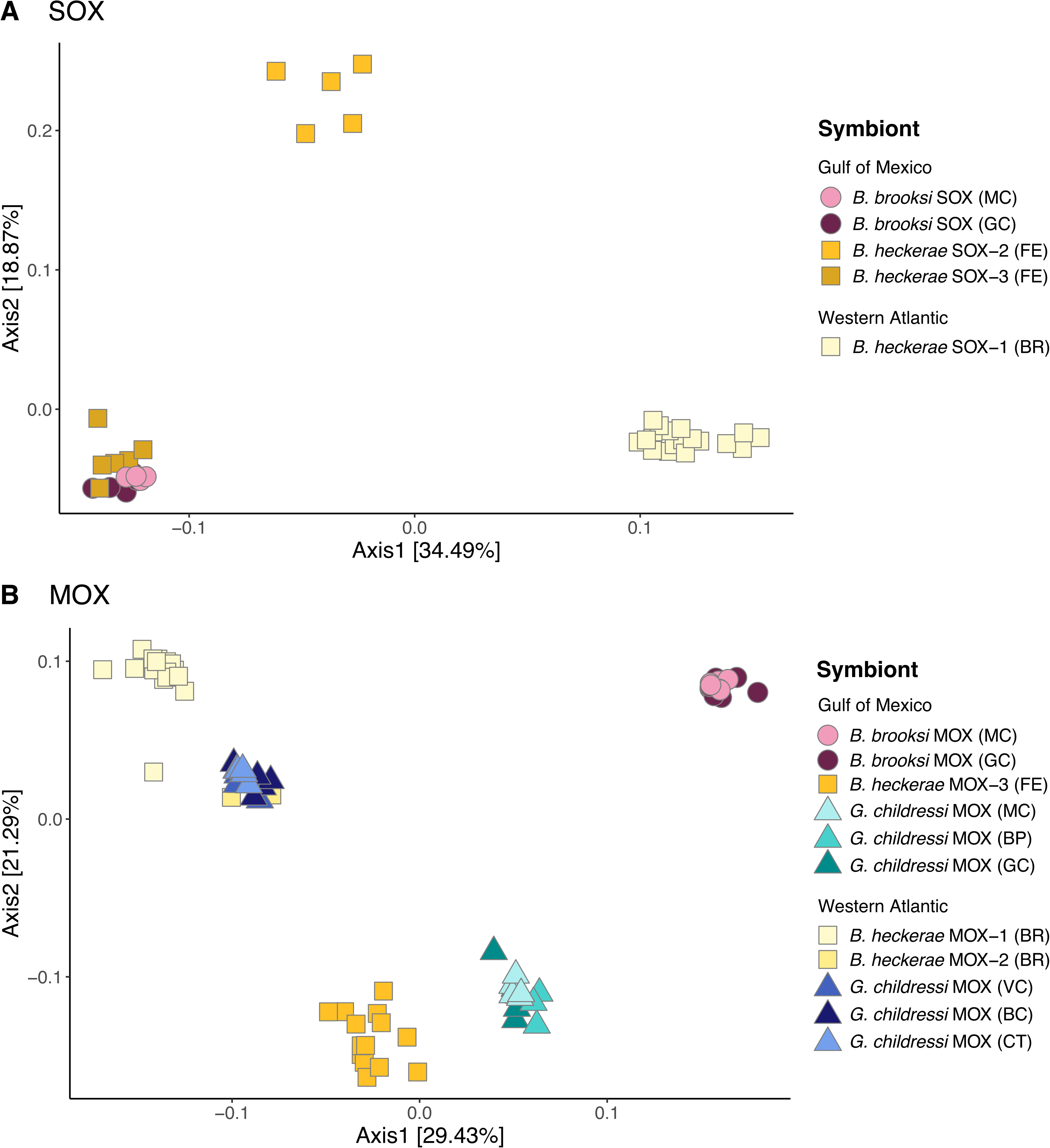
Principal component plots of KEGG enzyme presence/absence for (A) SOX symbiont species and (B) MOX symbiont species. Symbiont species showed both functional convergence and divergence based on geographic region and host species.

Comparative genomic analyses in the MOX symbionts identified five main clusters, consisting of (i) *B. brooksi* MOX, (ii) *B. heckerae* MOX-3 (GoM), (iii) *G. childressi* MOX (GoM), (iv) *B. heckerae* MOX-1 (WAM) and (v) *G. childressi* MOX (WAM) plus *B. heckerae* MOX-2 (WAM) (Fig. 4B). Overall, symbionts showed functional convergence by geographic region, with co-distributed symbiont species being functionally more alike than regional populations of the same symbiont species. For example, the *G. childressi* MOX populations in the WAM showed more functional similarities to the *B. heckerae* MOX-1 and MOX-2 symbiont species from that region than to the *G. childressi* MOX populations in the GoM. Notably, the *G. childressi* MOX and *B. heckerae* MOX-1 symbionts contained a soluble, iron-dependent methane monooxygenase in addition to the membrane-bound, copper-dependent methane monooxygenase observed in all other MOX symbionts and free-living relatives (Table S9, S10). Moreover, a NiFe hydrogenase and associated hydrogenase maturation and formation proteins were enriched in this symbiont cluster. By contrast, symbionts of different host species that co-occurred at the same locality, such as *G. childressi* and *B. brooksi* MOX from Mississippi Canyon and GC852, were divergent. In the *B. brooksi* MOX symbiont, specifically, multiple genes related to flagellar function and two-component systems for chemotactic behavior, biofilm formation and response to different environmental stimuli were overrepresented.

## Discussion

Although the patterns and processes of population differentiation have been studied in a handful of symbiotic associations^10,26–29^, research on both evolutionary convergence and divergence in these systems is rare. To start filling this gap, we assessed genetic and functional variation in three co-distributed mussel species that form single- or multi-partner symbioses with chemosynthetic bacteria at Atlantic cold seeps.

Our analyses indicate that the divide between the GoM and WAM likely represents a physical or ecological barrier to gene flow, as genetic differentiation was observed for both host and symbiont populations between these regions. These findings support observations in multiple other marine taxa^30–33^, where the Florida Peninsula has been noted as a genetic transition zone. Among host species, population structure was most pronounced in *B. heckerae*, which showed significant partitioning between Florida Escarpment and Blake Ridge based on nuclear and mitochondrial data.

The mitochondrial population structure reported here confirms a previous study based on COX1 barcodes, although the authors found that a shallower WAM *B. heckerae* population showed little mitochondrial divergence from GoM populations and that both regions should be well connected based on dispersal modelling results^34^. These findings imply that habitat differences or the deeper depths characterizing the Blake Ridge and Florida Escarpment seeps (>2000 m) drive population differentiation between these locations. For example, connectivity between these seeps might only occur through larval dispersal in slower bottom currents instead of the surface ocean. This could restrict gene flow among associated *B. heckerae* populations and might also explain why the shallower occurring *G. childressi*, which is known to disperse in surface waters^35^, was less differentiated among geographic regions. Alternatively, stronger population structure in *B. heckerae* compared to *G. childressi* could be due to symbiont-related ecological isolation, which has been hypothesized to drive host adaptive evolution in other species^36^. Interestingly, *B. heckerae* switched symbiont species between regions, while *G. childressi* retained the same symbiont species but associated with different local strains. This could be caused by contrasting dispersal capabilities among symbiont species, differences in the intensity of environmental pressures and/or host selection. Notably, F_ST_ outlier analyses identified multiple genes related to pattern recognition and immunological responses in *B. heckerae* that could play a role in host interactions with distinct symbiont species. Some outliers related to embryogenesis and germ cell development were also observed, suggesting that the two *B. heckerae* populations might be partly reproductively isolated. Although F_ST_ outliers were also detected in some reproductive and immunity genes in *G. childressi*, the locus-specific differentiation was lower, suggesting that barriers to gene flow are more permeable in this species. This raises the hypothesis that the strength of divergence at candidate symbiosis loci might dictate whether hosts associate with distinct strains of the same symbiont species or completely different symbiont phylotypes.

Within geographic regions, host populations were undifferentiated, whereas symbiont populations showed genetic variation across seep sites, similar to patterns described in mollusk symbioses from deep-sea hydrothermal vents^10,27,37^. Although population differentiation levels varied among symbiont species, F_ST_ outliers in all species were frequently linked to cell envelope metabolism and DNA recombination or transposition. This could suggest possible adaptations in symbiont interactions with the local environment, including bacteriophages and the mussel host. Many F_ST_ outliers were associated with transposases, which are known to alter gene structures and expression and promote acquisition of novel functions through horizontal gene transfer, thereby promoting rapid adaptation to environmental change^38^. For instance, transposases are thought to mediate adaptive capacities in free-living marine bacterial communities that are exposed to strong ecological fluctuations^39^. As fluid flow at cold seeps can oscillate significantly^40–42^, transposase-driven genomic plasticity might be beneficial for chemosynthetic symbionts that need to adjust to the changing availability of chemical reductants.

Across symbiont species, evidence for both functional divergence and convergence was observed that was linked to the geographic divide between the GoM and WAM. For example, the WAM populations of *G. childressi* MOX and *B. heckerae* MOX-1 (but not MOX-2; see Supplementary Discussion) both encoded a soluble, iron-dependent methane monooxygenase (sMMO) along with the standard membrane-bound, copper-dependent enzyme (pMMO). This is surprising since sMMO is typically absent in methanotrophic symbionts^43^. Based on phylogenetic evidence, sMMO was present in the ancestors of extant group I methanotrophs and was subsequently lost in multiple lineages^44^, explaining its patchy distribution across methanotrophic bacteria. Therefore, the presence of sMMO in the *G. childressi* MOX and *B. heckerae* MOX-1 species likely represents a case of independent gene retention. sMMO is expressed under low copper levels, while pMMO is only active when copper is available^45^. The fact that sMMO was only observed in WAM MOX symbionts indicates that WAM seeps, but not GoM seeps, are periodically copper limited. Although we lack chemical data for the studied localities, higher copper availability at the GoM seeps is also indirectly supported by the overrepresentation of copper detoxification genes in the *Ca.* Thiodubiliella symbionts of *B. brooksi* and *B. heckerae*. Unlike pMMO, sMMO has a broad substrate specificity and can act on diverse alkanes, alkenes, aromatic hydrocarbons and halogens apart from methane^46^, suggesting that the WAM *G. childressi* MOX and *B. heckerae* MOX-1 symbionts use multiple reductants for their chemosynthetic metabolism. This is further highlighted by the fact that both symbionts contained a NiFe hydrogenase that might enable supplemental use of H_2_ as energy source.

Contrary to the regional-scale metabolic convergence, MOX symbiont species that co-occurred at the same location diverged in multiple functions, indicating that release from competition through niche partitioning is likely relevant to symbiont variation in sympatric host species, as has been suggested for other chemosynthetic symbioses^15^. For example, the *B. brooksi* and *G. childressi* MOX symbionts from Mississippi Canyon and GC852 differed in gene content, with the *B. brooksi* symbiont being enriched in motility and chemotaxis genes. These genes might aid in the localization of optimal geochemical microhabitats and host individuals during the free-living stage^47–49^, possibly contributing to niche partitioning among mussel holobionts.

In the SOX symbionts, most species showed potential for anaerobic respiration using nitrogen compounds, although only the GoM *Ca*. Thiodubiliella symbionts encoded gene sets for complete denitrification. Multiple *Ca.* Thioglobus species are incomplete denitrifiers that use nitrate as electron acceptor for sulfur oxidation under hypoxic conditions^50,51^. Partial nitrate reduction in these taxa is probably a metabolic adaptation that skips energetic costs associated with the final denitrification steps. In the WAM *Ca.* Thioglobus SOX-1 the absence of nitrous oxide reductase might also be related to limited copper availability since this enzyme requires copper as cofactor^52^. The SOX symbionts further differed in the type of nitrate reductase used for respiration. While most *Ca.* Thiodubiliella symbionts and, in some cases, *Ca.* Thioglobus SOX-2 from the GoM encoded the membrane-bound *Nar*-type nitrate reductase, the WAM *Ca.* Thioglobus SOX-1 encoded the periplasmic *Nap*-type nitrate reductase. *Nar* and *Nap* nitrate reductases differ in their oxygen tolerance, with *Nar* usually functioning under anaerobic conditions and *Nap* operating under aerobic conditions, although exceptions exist^53–55^. These findings highlight regional differences in the use of electron acceptors for chemosynthesis between SOX symbionts, with GoM symbionts likely switching between oxygen and nitrate and WAM symbionts using both acceptors simultaneously, possibly due to higher O_2_ limitation.

Altogether, our results indicate that symbiont populations of cold-seep associated mussels evolve through both divergent and convergent mechanisms. We observed regional convergence in a key gene for methanotrophy across MOX symbionts from three distinct host species, while SOX symbionts showed regional divergence in dissimilatory nitrate reductase types, reinforcing that adaptation to environmental conditions is likely a strong driver of symbiont variation. We further observed divergence in symbiont function when species co-occurred at the same locality, suggesting that symbiont-driven niche differentiation may be important in locally sympatric holobionts. Among host species, *B. heckerae* was genetically most structured and switched symbiont species between oceanic regions, whereas *G. childressi* was less differentiated and retained the same symbiont species across locations. These patterns might be due to differences in dispersal potential, the strength of natural selection and/or host-symbiont interactions among holobionts. These results provide novel insights into the evolutionary processes shaping horizontally transmitted microbial symbioses.

## Materials and Methods

### Sample collection, DNA extraction and metagenomic sequencing

216 mussel samples were collected with ROVs, HOVs or camera-guided Smith-McIntyre grabs from eight cold seep sites at depths of 377–3287 m along the WAM and the GoM during research cruises between 2016 and 2022 (Fig. 1A; Table S1). On board the ship, gill tissues were dissected and stored at -80°C. Species identities were determined morphologically and confirmed through molecular barcoding of the cytochrome-c-oxidase subunit I (COX1) gene using bathymodiolin-specific primers^21^. DNA was extracted with the E.Z.N.A. Mollusc & Insect DNA kit (Omega Bio-tek, Norcross, GA, USA) and sent to Psomagen Inc. (Rockville, MA, USA) for library preparation and sequencing. Libraries were constructed with the Twist 96-Plex kit (Twist Bioscience, San Francisco, CA, USA) and 150 bp paired-end sequenced to a mean depth of 38,174,323 total reads on a NovaSeqX Plus instrument (Illumina, San Diego, CA, USA) (Table S1). The resulting raw reads were quality-checked with FastQC (https://github.com/s-andrews/FastQC) and trimmed with fastp^56^ and Trimmomatic^57^. Residual laboratory contamination was removed by mapping the trimmed reads against the human and PhiX genomic references with bowtie2^58^.

### Symbiont binning and phylogenetic analyses

To determine the taxonomic identity of sulfur-oxidizing and methane-oxidizing symbionts within each host individual the cleaned metagenomic reads of each sample were first assembled with metaSPAdes^59^ and then grouped into bins with metaBAT^60^ and MaxBin^61^ as described previously^37^. The best non-redundant MAG for each symbiont species and host individual was determined with DAS Tool^62^ and taxonomically delineated with GTDB-tk^63^ and fastANI^64^. MAGs were then quality-checked with CheckM^65^ based on either Methylococcales- or Gammaproteobacteria-specific marker genes. In general, MAGs with >85% completeness and <10% contamination were kept for further analysis (Table S2). However, because of the low amount of *B. heckerae* symbiont MAGs recovered with >85% completeness, for this host species we included additional MAGs that were at least 75% complete (Table S2). The relative abundance of the different SOX and MOX symbiont species within each host sample was determined with coverM^66^ based on the most complete representative MAG per symbiont species. To assess the phylogenetic relationships of the recovered symbiont species to other sulfur- and methane-oxidizing bacteria within the Thioglobaceae and Methylomonadaceae we constructed phylogenomic trees based on 33 and 297 single-copy core gene clusters (--mcl 2), respectively. Amino-acid based phylogenies were computed with IQ-Tree2^67^ in Anvi’o^68^ (parameters: -m MFP -B 1000 -bnni -alrt 1000 -T AUTO), using one representative MAG per symbiont species and location and related bacterial genomes from the National Center for Biotechnology Information for context (Table S11).

### Host population genetic analyses

To assess genetic variation within the three mussel host species, cleaned reads of each individual per species were mapped against high-quality reference genomes of *Gigantidas childressi* (GCA_965235005.1), *Bathymodiolus heckerae* (GCA_965117275.1) and *B. brooksi* (GCA_963680875.1) available through the Aquatic Symbiosis Genomics Project (https://portal.aquaticsymbiosisgenomics.org). Read alignments were adjusted by marking optical duplicates (https://broadinstitute.github.io/picard/), correcting alignments around indel regions and recalibrating base quality scores^69^. We subsequently inferred genotype likelihoods with ANGSD^70^ assuming Hardy-Weinberg equilibrium (HWE). Bi-allelic variant sites were filtered based on probability of being variable (-SNP_pval 1e-6), strand and heterozygote bias (-sb_pval 0.05 -hetbias_pval 0.05), deviation from HWE (-HWE_pval 0.05) and the quality of sequenced bases (-minQ 20) and read mappings (-minMapQ 30 -only_proper_pairs - remove_bads -unique_only). We further chose depth cutoffs based on the global read depth distribution to exclude potentially paralogous sites. Poorly covered genomic regions were ignored by requiring a site to be represented by a minimum of 50% of samples. Initial analyses in *G. childressi* indicated that inclusion of the full sample set resulted in relatively low amounts of SNPs (6,503). To improve SNP recovery in this species the sample set was therefore reduced to 15 samples per seep site for the computation of covariance matrices and overall population structure. For all other, population-specific analyses, such as SFS inference that formed the basis of pairwise F_ST_ calculations, all samples per population were used. Linkage disequilibrium between variant sites was calculated with Plink2^71^ in 20 kb windows using a step size of 2000 and a pairwise r^2^ threshold of 0.5. ANGSD analyses were then repeated considering only unlinked variants with the -sites and -rf options. As sample sizes were below 50 in both *B. heckerae* and *B. brooksi*, these analyses were only performed for *G. childressi*. However, patterns of population structure based on linked and unlinked sites in *G. childressi* were relatively similar (see below), confirming that inclusion of linked sites did not majorly impact the results. Pairwise F_ST_s between populations of each species were computed based on the final set of variant sites with ANGSD’s realSFS program. Potential candidate SNPs under positive selection were identified by selecting the 1% upper percentile of F_ST_ outliers. In the case of *G. childressi*, populations within the WAM and GoM were pooled for this analysis. The function of genes associated with the identified outlier SNPs was determined through homology information available for the different mussel reference genomes (https://ftp.ebi.ac.uk/pub/ensemblorganisms/).

As a complement to the population genetic analyses based on the nuclear genome, we assessed mitochondrial DNA variation across locations for each host species. Mitogenomes for each sample were assembled with MitoFinder^72^ and manually curated, standardized and, if necessary, circularized in Geneious Prime (https://www.geneious.com). For each host species, closed mitogenomes were aligned with progressiveMauve^73^, automatically trimmed with trimAl^74^ and a maximum-likelihood phylogeny was constructed with RAxML^75^ based on the GTRGAMMA model with 1000 bootstraps.

### Symbiont strain deconvolution and population genetic analyses

To infer the number and abundance of different strains within each symbiont species we used StrainFinder^76^ based on a reference set of 59 MOX and 93 SOX single-copy genes that we identified with Anvi’o^68^ (see below). Data were pre-processed as recommended in the StrainFinder documentation (https://github.com/cssmillie/StrainFinder) and strain deconvolution analyses were run for N = 2 to N = 10 strains with the following settings: --max_reps 100 --dtol 1 --ntol 3 --converge --n_keep 3 --force_update --merge_out --msg. The best model among runs was selected using the Bayesian Information Criterion. If the number of estimated strains for a symbiont species equaled 10, analyses were repeated for up to 15 strains. Slight adjustments to the code were made to ensure compatibility with Python 3. In particular, the numerical optimization based on the NLP solver of the OpenOpt framework was replaced with the scipy.optimize.minimize function. StrainFinder only estimates the relative abundance of strains within species but not across species. To obtain the actual abundance of each strain across samples and symbiont species we therefore multiplied the strain frequency with the species relative abundance obtained with coverM (see above). Heatmaps of strain abundances were produced with the complexheatmap package in R^77,78^ based on Bray-Curtis dissimilarities and average linkage.

As a complement to the strain inference analyses, we quantified population genetic differentiation for each symbiont species that was sampled across multiple locations. Species-specific pangenomes were generated from the final set of corresponding high-quality MAGs with Panaroo^79^ to provide references for population genomic analyses. Genetic variation within each symbiont species was assessed with FreeBayes^80^ following a previously published pipeline^10,27,37^ and using at least 17X of pangenome coverage per sample. Pairwise F_ST_s between symbiont populations at different seep sites were calculated with the scikit-allel Python package as described in Breusing et al. (2022)^10^. As in the host, potential variants under selection were identified by determining the 1% upper percentile of F_ST_ outliers or those with fixed differences. The function of genes containing these SNPs was assessed through Prokka annotations^81^ obtained during the pangenome reconstructions as well as complementary searches with blastp^82^ and eggNOG-mapper^83^.

### Symbiont comparative genomics and functional analyses

To identify functional differences and similarities among sulfur- and methane-oxidizing symbiont species across *Gigantidas* and *Bathymodiolus* hosts, we created Thioglobaceae and Methylomonadaceae specific pangenomes in Anvi’o^68^, using MCL clustering thresholds of 6 and 8, respectively. For *G. childressi*, the five best scoring MAGs for each seep location were used given that Anvi’o has a restriction of 100 total genomes for pangenome analyses. Presence/absence of KEGG enzymes was assessed with the “anvi-estimate-metabolism” program and visualized in a principal coordinate plot in R^78^. Functional enrichment among metabolically and genetically divergent symbiont groups across host species and locations was then computed with the “anvi-compute-functional-enrichment-in-pan” program using the KOfam database as annotation source. KOfam terms were considered enriched if their adjusted q value was equal or lower to 0.05.

## Data Availability

Raw sequence reads and MAG assemblies are available at the National Center for Biotechnology Information under BioProject PRJNA1443189.

## Ethics Statement

Ethical approval from an institutional review board was not required for this study.

## Acknowledgements

We thank the captains, crews, ROV and HOV pilots and the science parties of the R/V *Endeavor* (EN647), R/V *Atlantis* (AT36-01, AT50-04) and R/V *Thomas G. Thompson* (TN391) for supporting the sample collections that made this study possible. We also want to thank Russ Shomberg, Nick Chaloux, and Rendhy Sapiie for their work building and operating the camera-guided Smith-McIntyre grab used to collect mussels on EN647, Michelle Hauer, Erin Frates, and Ben Toles for their work on EN647 processing samples, Doreen McVeigh for collecting mussels on AT36-01, Abi Goodman for helping with DNA extractions and Psomagen Inc. for preparing and sequencing the metagenomic libraries for the analyzed mussel samples. The Unity HPC is gratefully acknowledged for providing computational resources and technical support for the bioinformatic analyses performed in this publication. This work was supported by National Science Foundation awards OCE-2141742 to RAB, OCE-1851286 to SMA, OCE-1851383 to CMY and OCE-1851421 to D. Eggleston, as well as a Rhode Island Endeavor Program shiptime award to RAB that funded EN647. Cruise AT36-01 was a UNOLS Early Career Training Program funded by NSF (OCE-1641453, OCE-1638805, OCE-1214335, OCE-1655587, and OCE-1649756) and ONR (N00014-15-1-2583).

## References

1. Brucker, R. M. & Bordenstein, S. R. Speciation by symbiosis. Trends Ecol. Evol. 27, 443–451 (2012).

2. Chow, J., Lee, S. M., Shen, Y., Khosravi, A. & Mazmanian, S. K. Host–Bacterial Symbiosis in Health and Disease. in Advances in Immunology vol. 107 243–274 (Elsevier, 2010).

3. McFall-Ngai, M. Symbiosis takes a front and center role in biology. PLoS Biol. 22, e3002571 (2024).

4. McFall-Ngai, M. et al. Animals in a bacterial world, a new imperative for the life sciences. Proc. Natl. Acad. Sci. 110, 3229–3236 (2013).

5. Rosenberg, E. & Zilber-Rosenberg, I. Microbes Drive Evolution of Animals and Plants: the Hologenome Concept. mBio 7, e01395–15 (2016).

6. Bright, M. & Bulgheresi, S. A complex journey: transmission of microbial symbionts. Nat. Rev. Microbiol. 8, 218–230 (2010).

7. Russell, S. L. Transmission mode is associated with environment type and taxa across bacteria-eukaryote symbioses: a systematic review and meta-analysis. FEMS Microbiol. Lett. 366, (2019).

8. Ganesan, R., Wierz, J. C., Kaltenpoth, M. & Flórez, L. V. How It All Begins: Bacterial Factors Mediating the Colonization of Invertebrate Hosts by Beneficial Symbionts. Microbiol. Mol. Biol. Rev. 86, e00126–21 (2022).

9. Bos, J. T., McManus, L. C., Ravago-Gotanco, R. & Pinsky, M. L. Contrasting patterns of seascape genetics in *Acropora* cf. tenuis and their symbiotic algae. Preprint at 10.64898/2026.04.07.716991 (2026).

10. Breusing, C., Genetti, M., Russell, S. L., Corbett-Detig, R. B. & Beinart, R. A. Horizontal transmission enables flexible associations with locally adapted symbiont strains in deep-sea hydrothermal vent symbioses. Proc. Natl. Acad. Sci. 119, e2115608119 (2022).

11. Elmer, K. R. & Meyer, A. Adaptation in the age of ecological genomics: insights from parallelism and convergence. Trends Ecol. Evol. 26, 298–306 (2011).

12. Yeaman, S. et al. Convergent local adaptation to climate in distantly related conifers. Science 353, 1431–1433 (2016).

13. Moreira, L. R. & Smith, B. T. Convergent genomic signatures of local adaptation across a continental-scale environmental gradient. Sci. Adv. 9, eadd0560 (2023).

14. Jackson, J. M. et al. Local adaptation across a complex bioclimatic landscape in two montane bumble bee species. Mol. Ecol. 29, 920–939 (2020).

15. Beinart, R. A. et al. Evidence for the role of endosymbionts in regional-scale habitat partitioning by hydrothermal vent symbioses. Proc. Natl. Acad. Sci. U. S. A. 109, E3241–3250 (2012).

16. Freeman, C. J. et al. Microbial symbionts and ecological divergence of Caribbean sponges: A new perspective on an ancient association. ISME J. 14, 1571–1583 (2020).

17. Iglesias-Prieto, R., Beltrán, V. H., LaJeunesse, T. C., Reyes-Bonilla, H. & Thomé, P. E. Different algal symbionts explain the vertical distribution of dominant reef corals in the eastern Pacific. Proc. R. Soc. Lond. B Biol. Sci. 271, 1757–1763 (2004).

18. Olu, K. et al. Biogeography and Potential Exchanges Among the Atlantic Equatorial Belt Cold-Seep Faunas. PLoS ONE 5, e11967 (2010).

19. Duperron, S. et al. Diversity, relative abundance and metabolic potential of bacterial endosymbionts in three *Bathymodiolus* mussel species from cold seeps in the Gulf of Mexico. Environ. Microbiol. 9, 1423–1438 (2007).

20. Faure, B., Schaeffer, S. W. & Fisher, C. R. Species Distribution and Population Connectivity of Deep-Sea Mussels at Hydrocarbon Seeps in the Gulf of Mexico. PLoS ONE 10, e0118460 (2015).

21. Olu-Le Roy, K., Cosel, R. V., Hourdez, S., Carney, S. L. & Jollivet, D. Amphi-Atlantic cold-seep *Bathymodiolus* species complexes across the equatorial belt. Deep Sea Res. Part Oceanogr. Res. Pap. 54, 1890–1911 (2007).

22. Coykendall, D. K. et al. Molecular characterization of *Bathymodiolus* mussels and gill symbionts associated with chemosynthetic habitats from the U.S. Atlantic margin. PLoS ONE 14, e0211616 (2019).

23. Raggi, L., Schubotz, F., Hinrichs, K. U., Dubilier, N. & Petersen, J. M. Bacterial symbionts of *Bathymodiolus* mussels and *Escarpia* tubeworms from Chapopote, an asphalt seep in the southern Gulf of Mexico. Environ. Microbiol. 15, 1969–1987 (2013).

24. Rubin-Blum, M. et al. Short-chain alkanes fuel mussel and sponge *Cycloclasticus* symbionts from deep-sea gas and oil seeps. Nat. Microbiol. 2, 17093 (2017).

25. Sogin, E. M., Leisch, N. & Dubilier, N. Chemosynthetic symbioses. Curr. Biol. CB 30, R1137–R1142 (2020).

26. Armstrong, K. C. et al. Fine Scale Geographic Variation of *Cladocopium* in *Acropora hyacinthus* Across the Palauan Archipelago. Ecol. Evol. 14, e70650 (2024).

27. Breusing, C., et al. Ecological differences among hydrothermal vent symbioses may drive contrasting patterns of symbiont population differentiation. mSystems 8, e0028423 (2023).

28. Cornwell, B. H. & Hernández, L. Genetic structure in the endosymbiont *Breviolum ‘muscatinei’* is correlated with geographical location, environment and host species. Proc. R. Soc. B Biol. Sci. 288, 20202896 (2021).

29. Perez, M., Angers, B., Young, C. R. & Juniper, S. K. Shining light on a deep-sea bacterial symbiont population structure with CRISPR. *Microb*. Genomics 7, (2021).

30. Anderson, J. D., O’Leary, S. J. & Cooper, P. T. Population Structure of Atlantic Croakers from the Gulf of Mexico: Evaluating a Single-Stock Hypothesis Using a Genomic Approach. Mar. Coast. Fish. 11, 3–16 (2019).

31. Portnoy, D. S. et al. Complex patterns of genetic population structure in the mouthbrooding marine catfish, Bagre marinus, in the Gulf of Mexico and U.S. Atlantic. Ecol. Evol. 14, e11514 (2024).

32. Black, K. L. et al. Evidence for gene flow from the Gulf of Mexico to the Atlantic Ocean in bonnethead sharks (*Sphyrna tiburo*). Ecol. Evol. 14, e70334 (2024).

33. Weinnig, A. M. et al. Population structure of *Desmophyllum pertusum* found along the United States eastern continental margin. BMC Res. Notes 17, 326 (2024).

34. Portanier, E. et al. Coupling large-spatial scale larval dispersal modelling with barcoding to refine the amphi-Atlantic connectivity hypothesis in deep-sea seep mussels. Front. Mar. Sci. 10, 1122124 (2023).

35. Arellano, S. M., Van Gaest, A. L., Johnson, S. B., Vrijenhoek, R. C. & Young, C. M. Larvae from deep-sea methane seeps disperse in surface waters. Proc. R. Soc. B Biol. Sci. 281, 20133276 (2014).

36. Osvatic, J. T. et al. Gene loss and symbiont switching during adaptation to the deep sea in a globally distributed symbiosis. ISME J. 17, 453–466 (2023).

37. Breusing, C. et al. Contrasting Genomic Responses of Hydrothermal Vent Animals and Their Symbionts to Population Decline After the Hunga Volcanic Eruption. Mol. Ecol. 34, e70126 (2025).

38. Casacuberta, E. & González, J. The impact of transposable elements in environmental adaptation. Mol. Ecol. 22, 1503–1517 (2013).

39. Vigil-Stenman, T., Ininbergs, K., Bergman, B. & Ekman, M. High abundance and expression of transposases in bacteria from the Baltic Sea. ISME J. 11, 2611–2623 (2017).

40. Cilenti, F., Oppo, D. & Macelloni, L. Controls on fluid discharge at cold seep-hydrate systems: 4D seismic monitoring of Woolsey Mound, Gulf of Mexico. Earth Planet. Sci. Lett. 648, 119087 (2024).

41. Sultan, N., Plaza-Faverola, A., Vadakkepuliyambatta, S., Buenz, S. & Knies, J. Impact of tides and sea-level on deep-sea Arctic methane emissions. Nat. Commun. 11, 5087 (2020).

42. Zhang, X. et al. Tidal modulation and temporal dynamics of methane emissions at the Formosa Ridge: insights from in-situ observations. Front. Mar. Sci. 12, 1633033 (2025).

43. Petersen, J. M. & Dubilier, N. Methanotrophic symbioses in marine invertebrates. Environ. Microbiol. Rep. 1, 319–335 (2009).

44. Osborne, C. D. & Haritos, V. S. Horizontal gene transfer of three co-inherited methane monooxygenase systems gave rise to methanotrophy in the Proteobacteria. Mol. Phylogenet. Evol. 129, 171–181 (2018).

45. Semrau, J. D., DiSpirito, A. A., Gu, W. & Yoon, S. Metals and Methanotrophy. Appl. Environ. Microbiol. 84, e02289–17 (2018).

46. Park, Y. R., Yoo, H. S., Song, M. Y., Lee, D.-H. & Lee, S. J. Biocatalytic Oxidations of Substrates through Soluble Methane Monooxygenase from *Methylosinus sporium* 5. Catalysts 8, 582 (2018).

47. Cambon-Bonavita, M.-A., Aubé, J., Cueff-Gauchard, V. & Reveillaud, J. Niche partitioning in the *Rimicaris exoculata* holobiont: the case of the first symbiotic Zetaproteobacteria. Microbiome 9, 87 (2021).

48. Colin, R., Ni, B., Laganenka, L. & Sourjik, V. Multiple functions of flagellar motility and chemotaxis in bacterial physiology. FEMS Microbiol. Rev. 45, fuab038 (2021).

49. Raina, J.-B., Fernandez, V., Lambert, B., Stocker, R. & Seymour, J. R. The role of microbial motility and chemotaxis in symbiosis. Nat. Rev. Microbiol. 17, 284–294 (2019).

50. Louca, S. et al. Integrating biogeochemistry with multiomic sequence information in a model oxygen minimum zone. Proc. Natl. Acad. Sci. 113, (2016).

51. Van Vliet, D. M. et al. The bacterial sulfur cycle in expanding dysoxic and euxinic marine waters. Environ. Microbiol. 23, 2834–2857 (2021).

52. Farrar, J. A., Zumft, W. G. & Thomson, A. J. Cu_A_ and Cu_Z_ are variants of the electron transfer center in nitrous oxide reductase. Proc. Natl. Acad. Sci. 95, 9891–9896 (1998).

53. Chen, X., Liu, C., Zhu, B., Wei, W. & Sheng, R. The Contribution of Nitrate Dissimilation to Nitrate Consumption in *narG*- and *napA*-Containing Nitrate Reducers with Various Oxygen and Nitrate Supplies. Microbiol. Spectr. 10, e00695–22 (2022).

54. Fischer, M., Falke, D., Pawlik, T. & Sawers, R. G. Oxygen-Dependent Control of Respiratory Nitrate Reduction in Mycelium of *Streptomyces coelicolor* A3(2). J. Bacteriol. 196, 4152–4162 (2014).

55. Luo, Y., Luo, L., Huang, X., Jiang, D. & Li, Z. narG, rather than napA, mediates aerobic nitrate reduction process in *Pseudomonas putida* Y-9. Water Res. X 29, 100437 (2025).

56. Chen, S., Zhou, Y., Chen, Y. & Gu, J. fastp: an ultra-fast all-in-one FASTQ preprocessor. Bioinforma. Oxf. Engl. 34, i884–i890 (2018).

57. Bolger, A. M., Lohse, M. & Usadel, B. Trimmomatic: a flexible trimmer for Illumina sequence data. Bioinformatics 30, 2114–2120 (2014).

58. Langmead, B. & Salzberg, S. L. Fast gapped-read alignment with Bowtie 2. Nat. Methods 9, 357–359 (2012).

59. Nurk, S., Meleshko, D., Korobeynikov, A. & Pevzner, P. A. metaSPAdes: a new versatile metagenomic assembler. Genome Res. 27, 824–834 (2017).

60. Kang, D. D. et al. MetaBAT 2: an adaptive binning algorithm for robust and efficient genome reconstruction from metagenome assemblies. PeerJ 7, e7359 (2019).

61. Wu, Y.-W., Simmons, B. A. & Singer, S. W. MaxBin 2.0: an automated binning algorithm to recover genomes from multiple metagenomic datasets. Bioinforma. Oxf. Engl. 32, 605–607 (2016).

62. Sieber, C. M. K. et al. Recovery of genomes from metagenomes via a dereplication, aggregation and scoring strategy. Nat. Microbiol. 3, 836–843 (2018).

63. Chaumeil, P.-A., Mussig, A. J., Hugenholtz, P. & Parks, D. H. GTDB-Tk: a toolkit to classify genomes with the Genome Taxonomy Database. Bioinforma. Oxf. Engl. 36, 1925–1927 (2019).

64. Jain, C., Rodriguez-R, L. M., Phillippy, A. M., Konstantinidis, K. T. & Aluru, S. High throughput ANI analysis of 90K prokaryotic genomes reveals clear species boundaries. Nat. Commun. 9, 5114 (2018).

65. Parks, D. H., Imelfort, M., Skennerton, C. T., Hugenholtz, P. & Tyson, G. W. CheckM: assessing the quality of microbial genomes recovered from isolates, single cells, and metagenomes. Genome Res. 25, 1043–1055 (2015).

66. Aroney, S. T. N. et al. CoverM: read alignment statistics for metagenomics. Bioinformatics 41, btaf147 (2025).

67. Minh, B. Q. et al. IQ-TREE 2: New Models and Efficient Methods for Phylogenetic Inference in the Genomic Era. Mol. Biol. Evol. 37, 1530–1534 (2020).

68. Eren, A. M. et al. Community-led, integrated, reproducible multi-omics with anvi’o. Nat. Microbiol. 6, 3–6 (2020).

69. Wilm, A. et al. LoFreq: a sequence-quality aware, ultra-sensitive variant caller for uncovering cell-population heterogeneity from high-throughput sequencing datasets. Nucleic Acids Res. 40, 11189–11201 (2012).

70. Korneliussen, T. S., Albrechtsen, A. & Nielsen, R. ANGSD: Analysis of Next Generation Sequencing Data. BMC Bioinformatics 15, 356 (2014).

71. Chang, C. C. et al. Second-generation PLINK: rising to the challenge of larger and richer datasets. Gigascience 4, s13742–015-0047–8 (2015).

72. Allio, R. et al. MitoFinder: Efficient automated large scale extraction of mitogenomic data in target enrichment phylogenomics. Mol. Ecol. Resour. 20, 892–905 (2020).

73. Darling, A. E., Mau, B. & Perna, N. T. progressiveMauve: Multiple Genome Alignment with Gene Gain, Loss and Rearrangement. PLoS ONE 5, e11147 (2010).

74. Capella-Gutiérrez, S., Silla-Martínez, J. M. & Gabaldón, T. trimAl: a tool for automated alignment trimming in large-scale phylogenetic analyses. Bioinformatics 25, 1972–1973 (2009).

75. Stamatakis, A. RAxML version 8: a tool for phylogenetic analysis and post-analysis of large phylogenies. Bioinformatics 30, 1312–1313 (2014).

76. Smillie, C. S. et al. Strain Tracking Reveals the Determinants of Bacterial Engraftment in the Human Gut Following Fecal Microbiota Transplantation. Cell Host Microbe 23, 229–240.e5 (2018).

77. Gu, Z., Eils, R. & Schlesner, M. Complex heatmaps reveal patterns and correlations in multidimensional genomic data. Bioinformatics 32, 2847–2849 (2016).

78. R Core Team. R: A Language and Environment for Statistical Computing. R Foundation for Statistical Computing (2023).

79. Tonkin-Hill, G. et al. Producing polished prokaryotic pangenomes with the Panaroo pipeline. Genome Biol. 21, 180 (2020).

80. Garrison, E. & Marth, G. Haplotype-based variant detection from short-read sequencing. Preprint at 10.48550/arXiv.1207.3907 (2012).

81. Seemann, T. Prokka: rapid prokaryotic genome annotation. Bioinformatics 30, 2068–2069 (2014).

82. Camacho, C. et al. BLAST+: architecture and applications. BMC Bioinformatics 10, 421 (2009).

83. Cantalapiedra, C. P., Hernández-Plaza, A., Letunic, I., Bork, P. & Huerta-Cepas, J. eggNOG-mapper v2: Functional Annotation, Orthology Assignments, and Domain Prediction at the Metagenomic Scale. Mol. Biol. Evol. 38, 5825–5829 (2021).

