## Supplementary Material for "Convergent and divergent patterns of holobiont evolution in co-distributed chemosynthetic symbioses"

### Supplementary Results

#### *Functional enrichment in SOX and MOX symbionts*

Functional enrichment analyses suggested that the identified symbiont metabolic groups differed significantly in multiple KOfam terms (Table S7). For example, the *Ca.* Thiodubiliella symbionts of *B. brooksi* and *B. heckerae* were enriched in proteins for copper detoxification, nitrous oxide reduction as well as some membrane transporters for metabolite import/export and the secretion of different proteins, including hemolysin and adhesin which play a role in host-symbiont interactions. In addition, particular restriction enzymes, proteases/peptidases and pilus formation proteins were overrepresented in these symbionts (Table S7). The *Ca.* Thiodubiliella symbionts appeared to be the only taxa able to perform complete denitrification of nitrate to molecular nitrogen using a membrane-bound *Nar*-type nitrate reductase, with subunit recovery being most complete in *B. brooksi*, possibly due to differences in binning efficiency (Table S8).

In the *B. heckerae Ca.* Thioglobus SOX-2 symbiont from Florida Escarpment overrepresented genes included various enzymes for the synthesis and modification of cell surface molecules, iron transport and storage and tellurium resistance. By contrast, *B. heckerae*'s *Ca.* Thioglobus SOX-1 symbiont from Blake Ridge showed enrichment in a periplasmic *Nap*-type nitrate reductase, a hydroxylamine dehydrogenase for the conversion of hydroxylamine to nitrate, a sulfide dehydrogenase involved in the generation of molecular sulfur stores, a multi-component $K^+$:$H^+$ transport system important for sodium tolerance and potassium transport, several cell envelope and surface related enzymes as well as genes involved in fatty acid degradation (Table S7).

The *G. childressi* MOX strains from the Gulf of Mexico were characterized by a CRISPR-Cascade complex, while the *B. heckerae* MOX-3 symbiont showed enrichment in assimilatory nitrate reductase, asparagine synthase and some enzymes involved in protein modification and antibiotic resistance. In the *B. brooksi* MOX symbiont overrepresented KOfam terms included multiple genes related to flagellar assembly and function as well as two-component system proteins linked to chemotactic behavior, biofilm formation and response to different environmental stimuli.

In the Western Atlantic, the *G. childressi* MOX and *B. heckerae* MOX-1 symbionts were distinguished by a soluble, iron-dependent broad-spectrum methane monooxygenase, which was present in addition to the typical membrane-bound, copper-dependent methane monooxygenase observed in all other MOX symbiont species and free-living relatives (Table S9, S10). Moreover, a NiFe hydrogenase and associated hydrogenase maturation and formation proteins were enriched in this symbiont cluster. Compared to the other symbionts, the Western Atlantic *G. childressi* MOX strains more frequently contained certain genes for the degradation of methanol, chitin and rare sugars, genes related to cell surface and envelope modification as well as particular genes involved in bacterial defense mechanisms and host interactions. In comparison, the *B. heckerae* MOX-1 symbiont exhibited enrichment in several type IV and VI secretion system proteins, signal transduction proteins and transcriptional regulators, genes related to anti-viral and antibiotic defense, some cell envelope and surface modification proteins as well as genes for the degradation of aromatic compounds, cholesterol and isonitriles. Lastly, the *B. heckerae* MOX-2 symbiont showed an overrepresentation of some transporters and enzymes involved in environmental stress response, including resistance to arsenite, mercury and antibiotics, as well as antioxidant and osmolytic defense. In addition, some genes for anaerobic energy metabolism, thiosulfate oxidation and energy generation from amino acid or fatty acid degradation were enriched in this symbiont.

### Supplementary Discussion

It is interesting to note that the *B. heckerae* MOX-2 symbiont from Blake Ridge did not contain sMMO. It is unlikely that this represents a binning artifact as MAGs of this symbiont species were nearly complete (87.99-94.63%), though the low number of genomes that could be obtained for analysis makes it difficult to draw general conclusions. However, since this symbiont was observed at relatively low sequence abundance in *B. heckerae* individuals relative to the dominant MOX-1 symbiont, it is probable that it performs complementary methane oxidation under different environmental conditions than MOX-1 or sustains the host in other ways. For instance, some genes for fatty acid catabolism were enriched in the *B. heckerae* MOX-2 symbiont, suggesting a mixotrophic lifestyle. Evidence for heterotrophic metabolism was also observed in other MOX symbionts from the Western Atlantic, such as potential for the degradation of chitin and rare sugars (*G. childressi* MOX) and cholesterol (*B. heckerae* MOX-1). Facultative heterotrophy appears to be relatively common in chemosynthetic symbionts[1] and might be an adaptation to carbon limitation or fluctuating reductant concentrations that would allow these symbioses to be sustained even when methane flux vanes.

### Supplementary References

1. Seah, B. K. B. *et al.* Sulfur-Oxidizing Symbionts without Canonical Genes for Autotrophic $CO_2$ Fixation. *mBio* **10**, e01112-19 (2019).

### Supplementary Figures

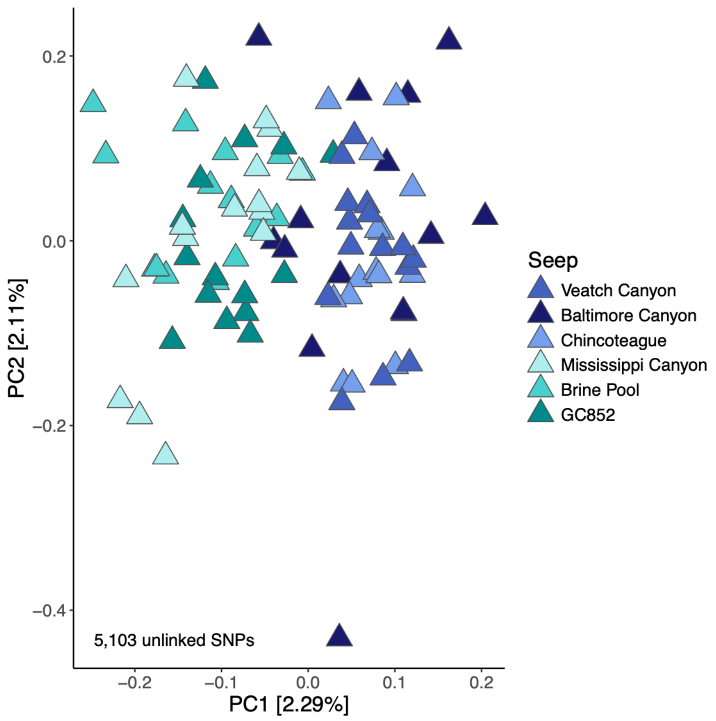


**Fig. S1** Principal component plot for *G. childressi* host populations based on unlinked genomic SNPs.

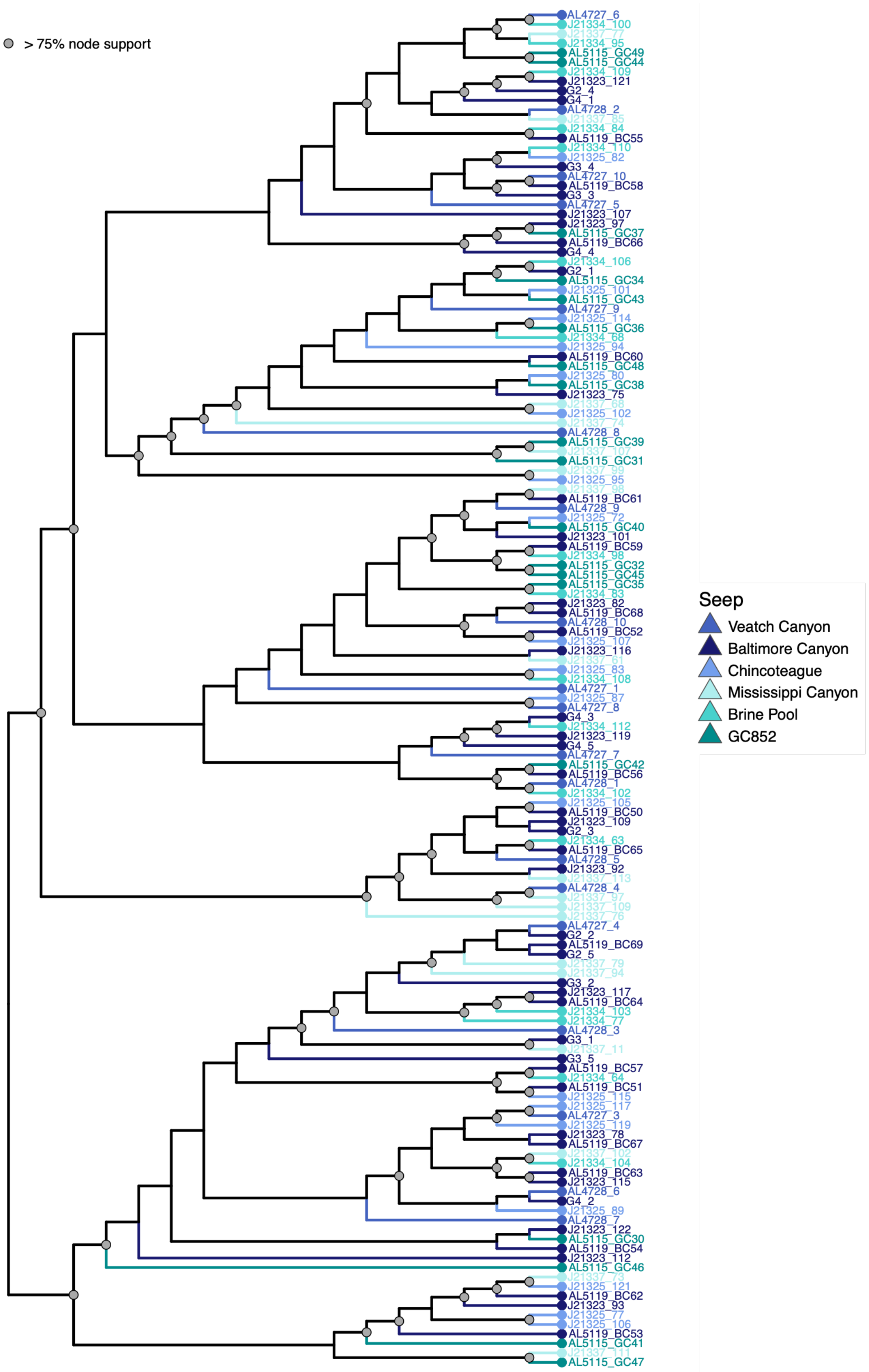


**Fig. S2** Cladogram of mitochondrial haplotypes for *G. childressi*.

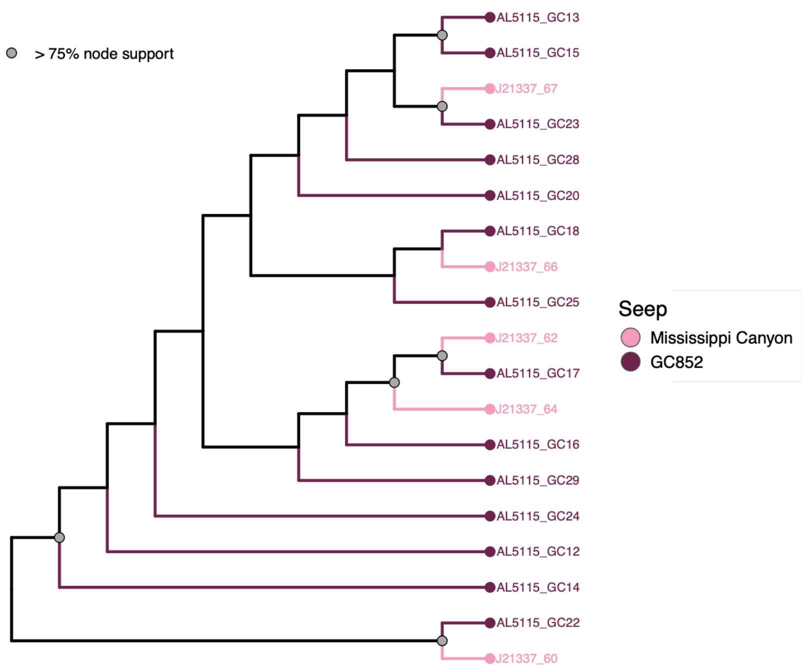


**Fig. S3** Cladogram of mitochondrial haplotypes for *B. brooksi*.

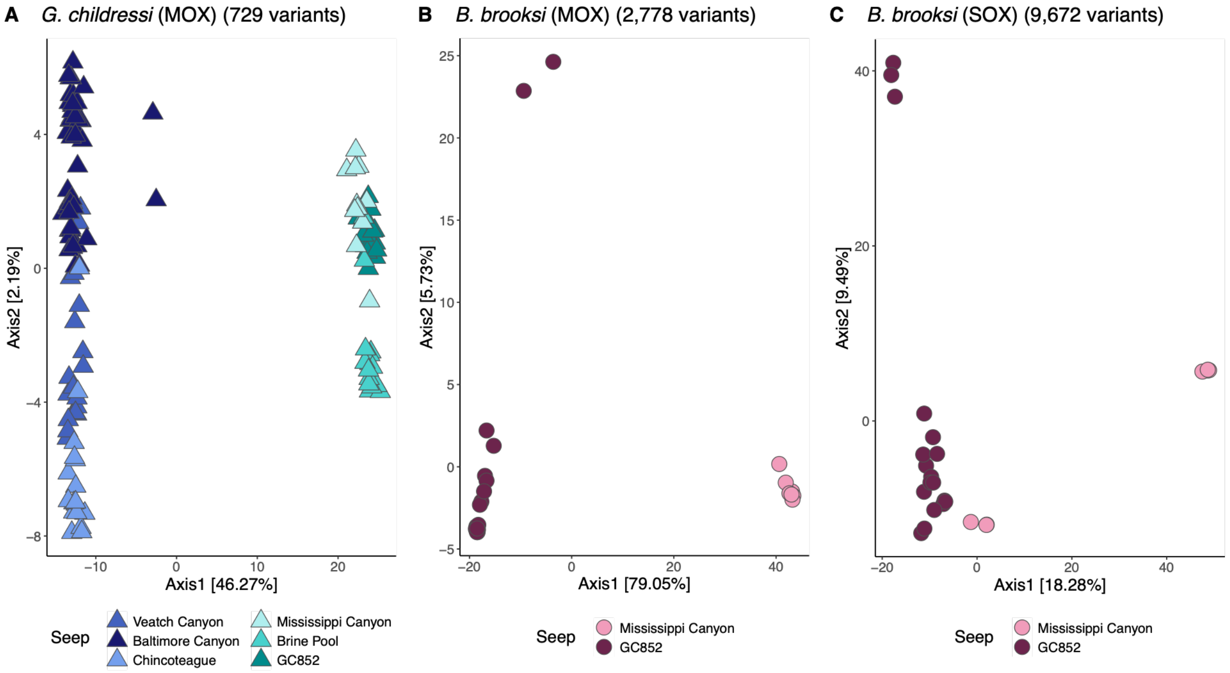


**Fig. S4** Principal component plots for symbiont populations of (A) *G. childressi* and (B, C) *B. brooksi* based on genomic variants. Populations of the *G. childressi* MOX symbiont were strongly partitioned between oceanic regions (Western Atlantic versus Gulf of Mexico), with lower albeit notable subdivision being observed within regions (A). In *B. brooksi*, populations of both the (B) MOX and (C) SOX symbiont species were structured between the Mississippi Canyon and GC852 seep sites within the Gulf of Mexico, though population differentiation was significantly higher in the MOX symbionts.
